# The L-alanosine gene cluster encodes a pathway for diazeniumdiolate biosynthesis

**DOI:** 10.1101/763607

**Authors:** Tai L. Ng, Monica E. McCallum, Christine R. Zheng, Jennifer X. Wang, Kelvin J. Y. Wu, Emily P. Balskus

## Abstract

*N*-nitroso-containing natural products are bioactive metabolites with antibacterial and anticancer properties. In particular, compounds containing the diazeniumdiolate (*N*-nitrosohydroxylamine) group display a wide range of bioactivities ranging from cytotoxicity to metal chelation. Despite the importance of this structural motif, knowledge of its biosynthesis is limited. Herein, we describe the discovery of a biosynthetic gene cluster in *Streptomyces alanosinicus* ATCC 15710 responsible for producing the diazeniumdiolate natural product L-alanosine. Gene disruption and stable isotope feeding experiments identified essential biosynthetic genes and revealed the nitrogen source of the *N*-nitroso group. Additional biochemical characterization of the biosynthetic enzymes revealed that the non-proteinogenic amino acid L-2,3-diaminopropionic acid (L-Dap) is synthesized and loaded onto a peptidyl carrier protein (PCP) domain in L-alanosine biosynthesis, which we propose may be a mechanism of handling unstable intermediates generated en route to the diazeniumdiolate. This research framework will facilitate efforts to determine the biochemistry of diazeniumdiolate formation.

---

*N*-nitroso-containing small molecules play prominent roles in human health, disease, and therapeutics development.^[1,2]^ In biological systems, nitric oxide synthases (NOSs) convert L-arginine into L-citrulline and nitric oxide (NO), which can undergo further oxidation to nitrite (NO_2_^−^) in aqueous systems. Both NO_2_^−^ and NO can react with secondary amines to afford *N*-nitrosamines, which are notorious environmental toxins and carcinogens, yet have been exploited as a bioactive pharmacophore in several *N*-nitrosourea-containing chemotherapeutic drugs.^[1,3]^ While many of these compounds are formed abiotically, the *N*-nitrosourea in streptozotocin (**2**) was recently reported to arise from the action of a metalloenzyme^[4]^ (SznF, Figure 1a), and the diazo functional group in cremeomycin (**4**)^[5]^ is thought to be produced via a transient *N*-nitrosamine intermediate (**3**) generated by an ATP-dependent enzyme (Figure 1b). Recent studies have linked bacterial diazeniumdiolate-containing metabolites to quorum sensing (**6**) and metal-acquisition (**7**), revealing an emerging ecological role for this group of natural products (Figure 1c).^[6,7]^ However, the genetic and biochemical basis for diazeniumdiolate biosynthesis remain poorly understood.

**Figure 1.**
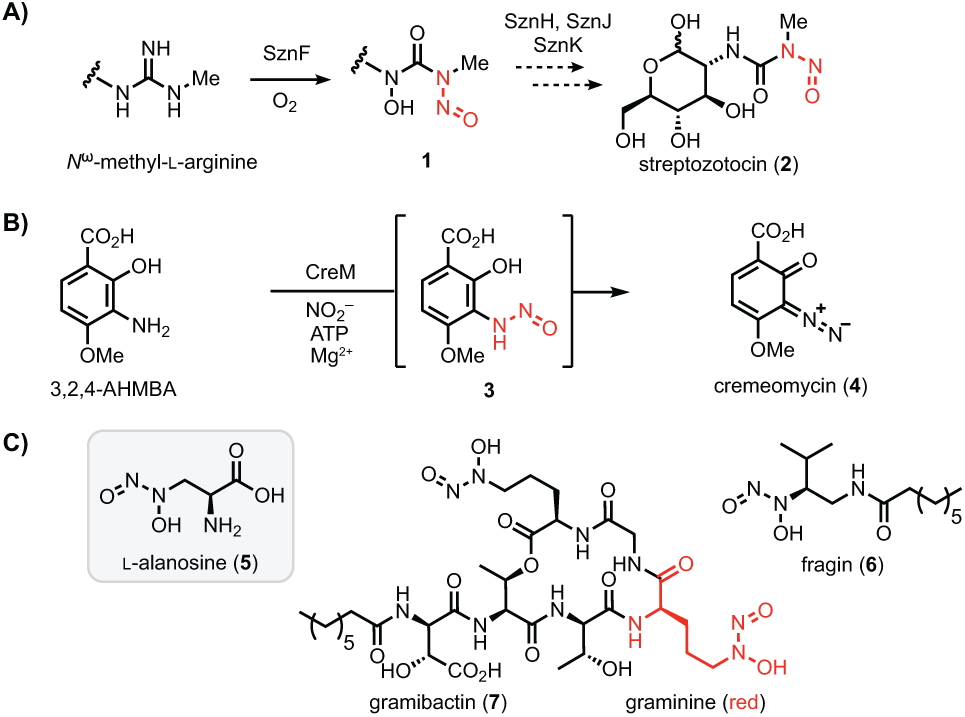
Oxidized nitrogen species in the biosynthesis of N–N bond-containing natural products. A) Biosynthesis of the *N*-nitrosourea group in streptozotocin. B) Biosynthesis of cremeomycin may proceed through an *N*-nitrosamine intermediate. C) Selected diazeniumdiolate-containing natural products.

L-alanosine (**5,** Figure 1c) is a naturally occurring diazeniumdiolate-containing amino acid produced by the bacterium *Streptomyces alanosinicus* ATCC 15710 that was discovered in the 1960s from a Brazilian soil sample.^[8]^ **5** has since been studied for its antibiotic, antiviral, and antitumor activities. The unusual diazeniumdiolate group confers metal chelating properties to **5**.^[9]^ This moiety also has been demonstrated to release NO upon metabolism of **5** by L-amino acid oxidases, generating reactive nitrogen species.^[10]^ In the decades following its isolation, **5** has been demonstrated to act as an antimetabolite targeting *de novo* purine biosynthesis and has been investigated in clinical trials to treat various tumors (SDX-102).^[11]^

Despite numerous studies exploring **5** as a potential therapeutic agent, the biosynthetic gene cluster responsible for its production has not been identified. Recently, the Eberl and Hertweck groups reported the biosynthetic gene clusters for the *N*-nitrosamines fragin (**6**)^[6]^ and gramibactin (**7**),^[7]^ revealing putative enzymes that could install the diazeniumdiolate moiety. HamC, which has been demonstrated to oxidize *p*-aminobenzoic acid to *p*-nitrobenzoic acid *in vitro*, is proposed to mediate N–N bond formation in fragin biosynthesis, but this putative reaction has yet to be verified *in vivo* and *in vitro*. Similarly, the SznF homolog GrbD from the gramibactin pathway is proposed to catalyze N–N bond formation, but its activity has not been demonstrated.^[6,12]^ Elucidating the biosynthesis of **5** would therefore improve our limited insights into enzymatic installation of the diazeniumdiolate, and more broadly the *N*-nitroso group. Here, we report the discovery of the L-alanosine (*ala*) biosynthetic gene cluster (Figure 2a). Using a combination of feeding studies, *in vivo* gene inactivation experiments, and *in vitro* biochemistry, we have revealed a plausible biosynthetic pathway, paving the way for further understanding of *N*-nitroso assembly in living organisms.

**Figure 2.**
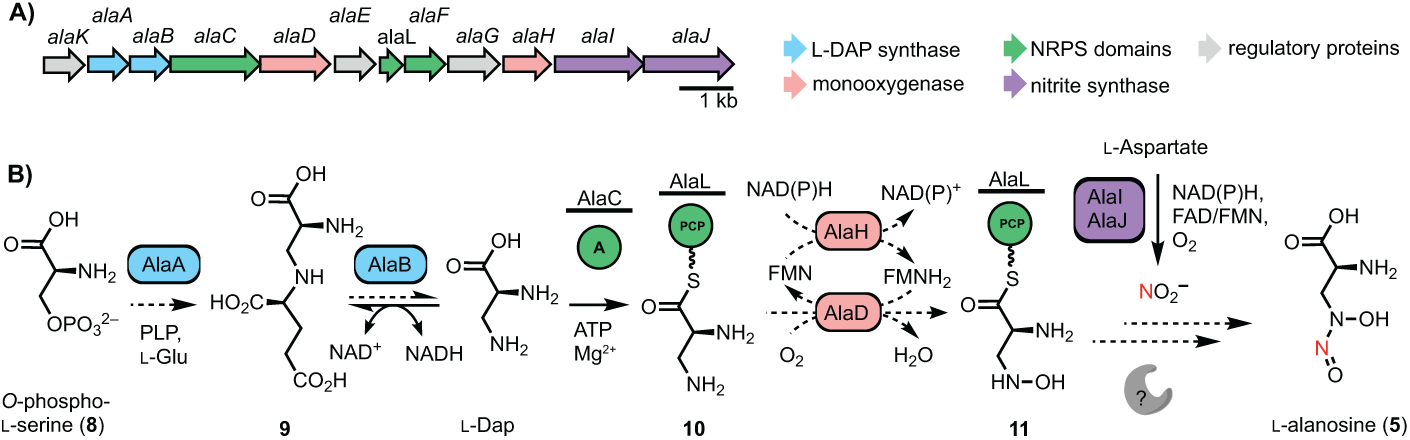
The L-alanosine gene cluster encodes a pathway for diazeniumdiolate biosynthesis. A) The putative L-alanosine (*ala*) gene cluster and B) biosynthetic hypothesis for the production of **5** from *O*-phospho-L-serine (**8**). Solid arrows represent enzyme reactions confirmed *in vitro*, dashed arrows represent proposed transformations. For detailed gene annotations see Table S1.

We initially hypothesized that **5** could be derived from *N*-hydroxylation and *N*-nitrosation of the putative biosynthetic precursor L-2,3-diaminopropionic acid (L-Dap, Figure 2b), a nonproteinogenic amino acid involved in several siderophore biosynthetic pathways.^[13-15]^ In staphyloferrin B biosynthesis, SbnA uses pyridoxal 5’-phosphate (PLP) as a cofactor to ligate *O*-phospho-L-serine (**8**) and L-glutamate to form a dipeptide *N*-(1-amino-1-carboxyl-2-ethyl)-glutamic acid (**9**) that is cleaved by deaminase SbnB to generate L-Dap (Figure S1).^[16]^ To identify the *ala* gene cluster, we sequenced the genome of *S. alanosinicus* ATCC 15710 and searched for homologs of the L-Dap biosynthetic genes. This strategy revealed a 13.5 kb genomic region that encodes homologs of SbnAB (AlaAB), a free-standing adenylation (A) domain (AlaC), a free-standing peptidyl carrier protein (PCP) domain (AlaL), and a thioesterase (TE) domain (AlaF) (Figure 2a, Table S1). These biosynthetic enzymes are encoded alongside homologs of the enzymes CreD (AlaJ) and CreE (AlaI), which are known to produce NO_2_^−^ from L-aspartic acid in cremeomycin biosynthesis,^[5]^ a predicted flavin-dependent *N*-hydroxylase (AlaD), and a potential flavin reductase (AlaH) (Figure 2a). The putative *ala* gene cluster also contains a transcriptional regulator (AlaK), a GAF-domain containing protein (AlaE) and a PAS-domain containing protein (AlaG) that likely regulate the transcription of this gene cluster.

Based on the predicted functions of these enzymes, we proposed a biosynthetic hypothesis for the assembly of **5** (Figure 2b). AlaAB would generate L-Dap, which could be loaded onto the phosphopantethienyl arm of the freestanding PCP AlaL by A domain AlaC. This biosynthetic logic parallels the proposed pathway for fragin construction in which diazeniumdiolate installation could occur on enzyme-tethered intermediates.^[6]^ The predicted flavin-binding enzyme AlaD and the flavin reductase AlaH could be a two-component flavin *N*-monooxygenase involved in forming *N*-hydroxy-L-Dap (**11**), and an unknown enzyme (potentially one of the remaining proteins encoded by the *ala* gene cluster) could install the *N*-nitroso group using NO_2_^−^ generated by AlaIJ. Notably, the *S. alanosinicus* genome does not encode homologs of any biosynthetic enzymes from the fragin and L-graminine pathways. Furthermore, the genome lacks homologs of SznF and KtzT, recently reported N–N bond-forming enzymes in streptozotocin and piperazate biosynthesis, respectively.^[4,17]^ Moreover, homologs of other suspected N–N bond-forming enzymes, Spb40/Tri28 from the s56-p1 and triacsin biosynthetic pathways^[18,19]^ and FzmP/KinJ from fosfazinomycin and kinamycin biosynthetic pathways, are also absent.^[20]^ Our bioinformatics analysis therefore suggests that biosynthesis of **5**

either employs a novel N–N bond-forming enzyme or generates the diazeniumdiolate functional group non-enzymatically.

To establish the link between the *ala* gene cluster and biosynthesis of **5**, we performed several gene inactivation experiments in *S. alanosinicus*. Production of **5** was abolished when *alaC, alaD*, and *alaI* were deleted via the well-established PCR-targeting and λ-red-mediated recombination platform (Figure 3a). This confirms that the activity of the A domain (AlaC), redox chemistry (via AlaD) and NO_2_^−^ generation (via AlaI) are all essential. When the *ΔalaI* mutant was supplemented with NO_2_^−^, **5** was not detected (Figure 3a, maroon line). This suggests that *N*-nitrosation of an L-alanosine precursor may be enzyme-mediated and may not occur spontaneously. Lastly, the deletion of *alaC* and concomitant loss of **5** imply that the NRPS machinery is necessary for biosynthesis and may facilitate the generation of an unstable, enzyme-bound intermediate during diazeniumdiolate assembly. Taken together, these gene-inactivation experiments confirmed the role of the *ala* gene cluster in the biosynthesis of **5** and identified several indispensable enzymes for further *in vitro* characterization.

**Figure 3.**
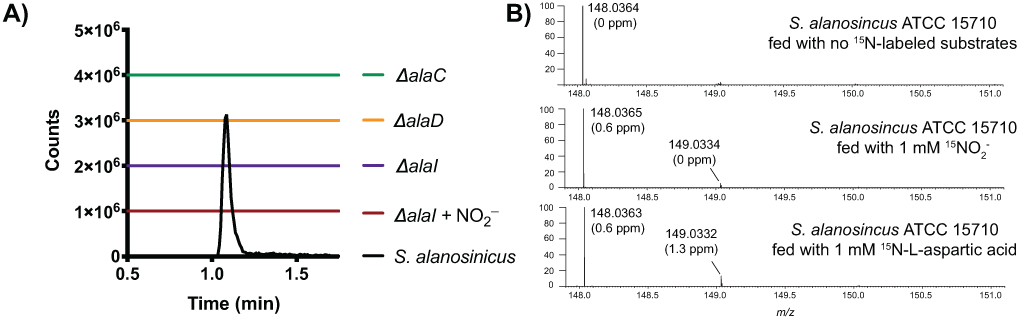
Gene inactivation and feeding studies link *ala* biosynthetic genes to L-alanosine (**5**) production A) EICs of **5** ([M–H]^−^ = 148.0364) in fermentation extracts of gene deletion mutants and wild type *S. alanosinicus*. The EICs are generated within a 5 ppm window B) Feeding studies using ^15^N sources with wild type *S. alanosinicus* to determine the source of the distal *N*-nitroso nitrogen atom of **5**. LC-MS/MS analysis confirmed the distal *N*-nitroso nitrogen was labeled when ^15^N-nitrite was fed (Figure S2). Higher incorporation of ^15^N into **5** was observed when 3 mM ^15^NO_2_^−^ was fed, although the cell viability was lower due to toxicity (Figure S3).

Because the putative *ala* gene cluster encodes homologs of the nitrite-generating enzymes CreD and CreE (AlaJ and AlaI, respectively, Figure 4a), we proposed that the precursor of the distal nitrogen of the *N*-nitroso group is NO_2_^−^. In addition to its role in non-enzymatic *N*-nitrosation reactions, NO_2_^−^ is used biosynthetically as a source of nitrogen in the formation of other N–N bond linkages, including diazo and hydrazide groups.^[20,21]^ To test this hypothesis, we overexpressed and purified N-His_6_-AlaI and N-Strep-AlaJ, and we demonstrated their ability to generate NO_2_^−^ from L-aspartic acid with the necessary cofactors (Figure 4b). We next fed ^15^NO_2_^−^ to the fermentation cultures of *S. alanosinicus* and observed incorporation of ^15^N into **5** (∼ 5%, Figure 3b). Tandem high resolution-mass spectrometry (HR-LCMS/MS) revealed the distal *N*-nitroso nitrogen is labeled (Figure S2). Feeding ^15^NO_3_^−^ also resulted in lower enrichment of ^15^N-labeled **5** (Figure S4); this could potentially arise from conversion of NO_3_^−^ to NO_2_^−^ by the nitrate reductases encoded in *S. alanosinicus*. Finally, we also fed ^15^N-L-aspartic acid, the presumed precursor of ^15^NO_2_^−^ and observed enrichment of ^15^N-**5** (∼ 10%, Figure 3b). Together, these results suggest that the biosynthesis of **5** employs NO_2_^−^ generated from L-aspartic acid, and the distal nitrogen of the diazeniumdiolate is derived from NO_2_^−^. Given that the aforementioned supplementation of the *ΔalaI* strain with free NO_2_^−^ did not lead to detectable production of **5**, incorporation of NO_2_^−^ may require interaction of AlaIJ with a putative N–N bond-forming enzyme or a PCP-tethered substrate.

**Figure 4.**
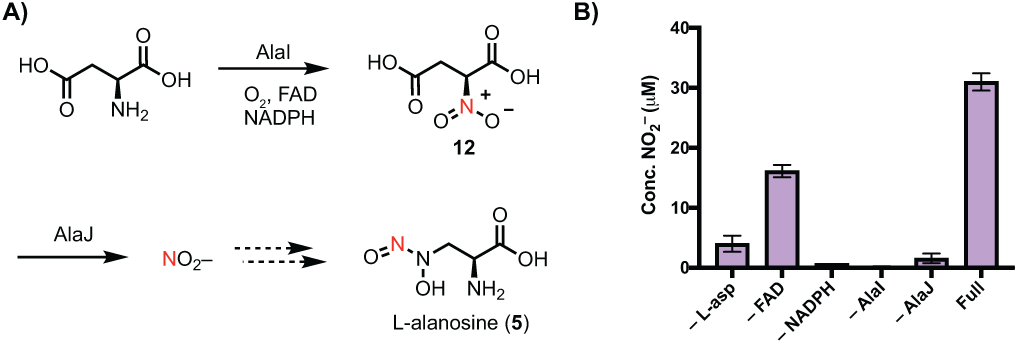
AlaI and AlaJ are a nitrite synthase. A) Proposed mechanism of NO_2_^−^ generation from L-aspartic acid by flavin dependent monooxygenase AlaI and nitrosuccinate lyase AlaJ B) NO_2_^-^ production detected by the Griess assay of FAD (10 μM), NADPH (5 mM), N-Strep-AlaJ (5 μM), and N-His_6_-AlaI (5 μM) and L-aspartic acid (1 mM) in reaction buffer (50 mM HEPES, 10 mM MgCl_2_, pH = 8.0) after a 1 h incubation at room temperature. AlaI and AlaJ produce NO_2_^−^ from L-aspartic acid *in vitro.* AlaI purified with FAD, hence we still observed activity without exogenous FAD added. Data are mean ± standard deviation (s.d.) of two technical replicates.

Having established the origin of the *N-*nitroso group in **5**, we next sought to confirm the role of L-Dap in this biosynthetic pathway. We first overexpressed and purified *N*-His_6_-AlaB for *in vitro* biochemical assays. SbnB, a homolog of AlaB, was previously characterized by performing the L-Dap synthase reaction in reverse to generate **9** (Figure S1).^[16]^ We employed the same strategy to characterize AlaB by incubating the enzyme with L-Dap and NADH. We observed rapid consumption of NADH and production of **9** (Figure 2b, Figure S5 and S6). Therefore, AlaB is a functional homolog of the previously characterized L-Dap synthase component SbnB.^[16]^

The involvement of a free-standing A domain in the biosynthesis of **5** and its putative loading of L-Dap are unusual, as there are only three examples of biochemically characterized A domains that utilize this amino acid.^[22-24]^ A domain active sites contain 8-10 binding-pocket residues that have been found to be predictive of substrate specificity.^[25]^ Several bioinformatic methods that predict of A domain substrate specificity from these binding-pocket residues have been developed.^[26]^ However, these tools^[27]^ failed to predict, L-Dap as the substrate of AlaC. The same is true of other A domains that have been experimentally demonstrated to activate L-Dap (Table S1).

To test if AlaC can recognize and load L-Dap, we overexpressed and purified N-His_6_-tagged AlaC and N-His_6_-C-His_6_-tagged AlaL for *in vitro* biochemical assays. The preferred substrate of AlaC was confirmed to be L-Dap using the ATP-[^32^P]Pi exchange assay (Figure 5a). The ability of purified apo-AlaL to undergo successful posttranslational modification was determined via incubation with the promiscuous phosphopantetheinyl (ppant) transferase Sfp and the fluorescent coenzyme A (CoA) analog BODIPY-CoA.^[28]^ Finally, we demonstrated that isotopically labeled L-^15^N_2_-Dap was activated by AlaC and loaded onto the ppant arm of holo-AlaL using whole-protein mass spectrometry (Figure 5b). Thus, the NRPS machinery encoded in the *ala* gene cluster is capable of activating L-Dap and loading it onto the PCP AlaL. Given that *alaC* was determined to be necessary for the production of **5**, the AlaL-tethered L-Dap aminoacyl thioester **10** (Figure 2b) is likely a biosynthetic intermediate.

**Figure 5.**
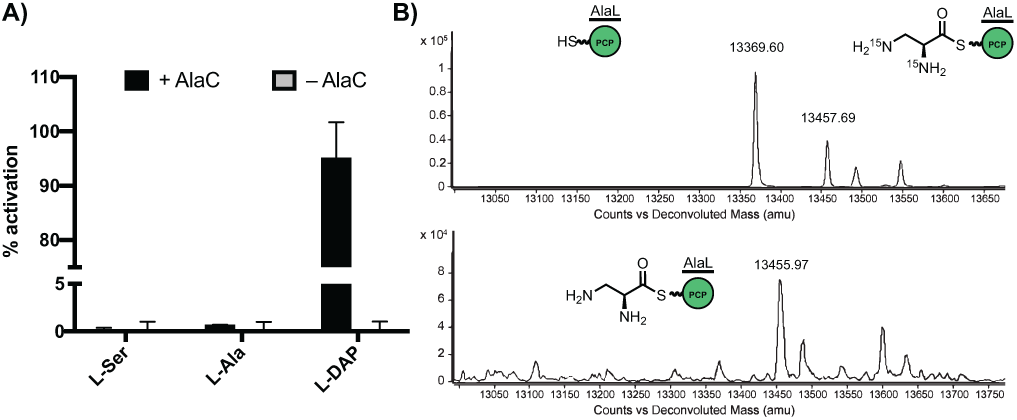
AlaC selectively activated L-Dap for loading onto AlaL. A) ATP-^32^PP_i_ exchange assay shows that AlaC preferentially activates L-Dap over other amino acids from 100 μL incubations of N-His_6_-AlaC (1 μM) with 5 mM dithiothreitol, 5 mM ATP, 1 mM of amino acid substrate, and 4 mM Na_4_PPi/[^32^P]PPi in reaction buffer (50 mM HEPES, 200 mM NaCl, 10 mM MgCl_2_, pH = 8) at room temperature for 30 min. Data are mean ± standard deviation (s.d.) of two biological replicates. B) Deconvoluted whole protein mass spectra (ESI+) showing holo-N-His_6_-AlaL-C-His_6_ (13369.60) is loaded with ^15^N_2_-L-Dap (13457.69) or L-Dap (13455.97). 50 μL reaction mixtures were set up with coenzyme A (1 mM), Sfp (5 μM), and N-His_6_-AlaL-C-His_6_ (20 μM) in a solution of reaction buffer (50 mM HEPES, 200 mM NaCl, 10 mM MgCl_2_, pH = 8). After incubation at room temperature for 2 h, AlaC (20 μM) and L-Dap or L-^15^N_2_-Dap (250 μM) were added, followed by ATP (5 mM) to initiate the reaction. Incubated for 1 h at room temperature.

In summary, we have identified a set of genes that are required for the biosynthesis of **5** in *S. alanosinicus*. We have demonstrated the importance of the freestanding NRPS biosynthetic machinery, both by generating genetic knockouts of *S. alanosinicus* and through *in vitro* biochemical assays. We have confirmed the role of L-Dap as a biosynthetic precursor by showing that AlaB is a functional homolog of the L-Dap synthase SbnB and characterizing a new L-Dap specific A domain. The previously characterized biosynthetic uses of L-Dap have been as a building block for NRPS natural products, so its utilization as the core of a small molecule is unusual. This may suggest that as yet undiscovered NRPS natural products contain L-alanosine building blocks in the same manner as gramibactin contains L-graminine building blocks.^[7]^ We have also demonstrated a critical role for NO_2_^−^ synthesis in this pathway, having confirmed through stable isotope feeding studies that L-aspartic acid the source of the *N*-nitroso group and shown that AlaI and AlaJ generate NO_2_^−^ from L-aspartic acid *in vitro*. The presence of known NO_2_^−^ generation enzymes in the *ala* gene cluster, and confirmation of their necessity for the production of **5**, provides preliminary insights into diazeniumdiolate formation. Coupled with a lack of known homologs of *N*–nitrosating enzymes in the *ala* gene cluster, these findings hint at a potentially distinct mechanism of N–N bond-formation in L-alanosine biosynthesis.

## Experimental Section

Complete experimental details are provided in the Supporting Information.

## Acknowledgements

We thank Dr. L. Zha for assistance with the ATP-PPi exchange assay, Dr. B. A. Schneider for assistance with the BODIPY-CoA loading assay, and Dr. A. J. Waldman for guidance with the nitrite production assay. M.E.M. acknowledges the Damon Runyon Foundation for a Postdoctoral Fellowship (DRG 2307-17). We acknowledge financial support from the NIH (DP2 GM105434), a Cottrell Scholar Award (to E.P.B.), a Camille Dreyfus Teacher-Scholar Award (to E.P.B.), and Harvard University.

