## Supporting Information for "The L-alanosine gene cluster encodes a pathway for diazeniumdiolate biosynthesis"

### 1. Materials and general methods

Oligonucleotide primers were synthesized by Integrated DNA Technologies (Coralville, IA) and Sigma-Aldrich (Atlanta, GA). Recombinant plasmid DNA was purified with a Qiaprep Kit from Qiagen (Germantown, MD). Gel extraction of DNA fragments and restriction endonuclease clean up were performed using Zymoclean™ Gel DNA Recovery Kit and DNA Clean & Concentrator kit from Zymo Research (Irvine, CA). DNA sequencing was performed by Genewiz (Boston, MA) and Eton Bioscience (Charleston, MA). Gibson assembly was conducted using New England Biolabs (NEB) (Ipswich, MA) Gibson Assembly® Master Mix following the manufacturer's protocol. Nickel-nitrilotriacetic acid agarose (Ni-NTA) resin was purchased from Qiagen and Thermo Scientific (Waltham, MA). Strep-Tactin® Sepharose® was purchased as a 50% suspension from IBA GmbH. SDS-PAGE gels were purchased from BioRad (Hercules, CA). Protein concentrations were determined by measuring absorption at 280 nm and using ExPASy ProtParam (http://web.expasy.org/protparam/) to calculate the extinction coefficients. Optical densities of *E. coli* cultures were determined with a DU 730 Life Sciences UV/Vis spectrophotometer (Beckman Coulter) by measuring absorbance at 600 nm. All water used experimentally, unless otherwise stated, was purified using a MilliQ (mQ) water purification system.

High-resolution mass spectral data for the synthetic compounds were obtained on a Bruker MicroQTOF-QII mass spectrometer fitted with a dual-spray electrospray ionization (ESI) source. The capillary voltage was set to 4.5 kV and the end plate offset to –500 V, the drying gas temperature was maintained at 190 ºC with a flow rate of 8 L/min and a nebulizer pressure of 21.8 psi. The liquid chromatography was performed using an Agilent Technologies 1100 series LC. Isopropanol, methanol, and water used for LC-ESI-MS were B & J Brand High Purity Solvents (Honeywell Burdick & Jackson).

### 2. Cultivation of *Streptomyces alanosinicus* ATCC 15710 and detection of L-alanosine (5).

*Streptomyces alanosinicus* ATCC 15710 was obtained from the Agricultural Research Service (ARS) Culture Collection. The organism was grown on mannitol-soy (MS) agar plates (20 g/L soy flour, 20 g/L mannitol, and 20 g/mL Bacto™ agar (Difco™)) at 30 ºC for five days. The spores were scraped and inoculated into 5 mL of International Streptomyces Project Medium 2 (ISP2) and allowed to shake at 30 ºC until saturation. 3 mL of this starter culture was inoculated into 100 mL production medium (50 g/L glucose, 10 g/L meat extract, 5 g/L $CaCO_3$, 1 g/L $(NH_4)_2SO_4$ 1 g/L $MgSO_4$, 1 mL of trace mineral solution) in a 250 mL baffled flask. The trace mineral solution consists of 0.5% (*w/v*) $CuSO_4$, 0.1% (*w/v*) $FeSO_4$, 0.2% (*w/v*) $ZnSO_4$, and 0.8% (*w/v*) $MnSO_4$. The fermentation was carried out for 5 days at 30 ºC shaking at 200 rpm. For detection of **5**, 750 µL of the culture was centrifuged at 13,000 rpm to remove cells, and the supernatant was added to an equivalent volume of acetone to further precipitate proteins and medium components. After vortexing, the insoluble material was removed by centrifugation at 13,000 rpm, and the sample was analyzed with LC-HRMS.

Detection of **5** and various compounds by LC-MS was carried out on a Thermo Scientific Dionex UltiMate 3000 UHPLC coupled to a Thermo Q Exactive Plus mass spectrometer system (Thermo Fisher Scientific Inc, Waltham, MA) equipped with an HESI-II electrospray ionization (ESI) source. Data were acquired with Chromeleon Xpress software for UHPLC and Thermo Xcalibur software version 3.0.63 for mass spectrometry and processed with Thermo Xcalibur Qual Browser software version 4.0.27.19. 8 μL of sample was injected onto the UHPLC including an HPG-3400RS binary pump with a built-in vacuum degasser and a thermostated WPS-3000TRS high performance autosampler. A Symmetry Shield RP 18 analytical column (2.1x150 mm, 3.5 μm) from Waters Corporation (Milford, MA) were used at the flow rate of 0.3 mL/min using 0.1% formic acid in water as mobile phase A and 0.1% formic acid in acetonitrile as mobile phase B. The column temperature was maintained at room temperature. The following gradient was applied: 0–2.4 min: 0% B isocratic, 2.4–2.5 min: 0–100% B, 2.5–6.1 min: 100% B isocratic, 6.1–6.2 min: 100–0% B, 6.2–12.2 min: 0% B isocratic. The MS conditions were as follows: negative ionization mode; full scan mass range, *m/z* 50 to 750; resolution, 140,000; AGC target, 1e6; maximum IT, 480 ms; PRM NCE, 20 for m/z 149.0334; resolution, 140,000; AGC target, 1e6; maximum IT, 480 ms; spray voltage, 3500 V; capillary temperature, 280 ˚C; sheath gas, 47.5; Aux gas, 11.25; probe heater temperature, 412.5 ˚C; S-Lens RF level, 50.00. A mass window of ±5 ppm was used to extract the ion of $[M–H]^–$ for the compound. The target was considered detected when the mass accuracy was less than 5 ppm and there was a match of isotopic pattern between the observed and the theoretical one and a match of retention time between those in real sample and standard.

### 3. Genome sequencing of *Streptomyces alanosinicus* ATCC 15710 and genomic DNA library construction

Genomic DNA was purified using the UltraClean® Microbial DNA Isolation Kit (Mo Bio). Library construction from genomic DNA, sequencing, and assembly were performed by Era7 Bioinformatics (St. Louis, MO). Next-generation sequencing used HiSeq2000 of two short-insert paired-end libraries (100 bp insert). Assembly of the reads resulted in a 10.3 MB of non-redundant sequence distributed over 236 contigs. Annotations were carried out using Era7's BG7.[1] The assembled data were converted into a local BLAST database using Geneious. The *Streptomyces alanosinicus* ATCC 15710 fosmid library was prepared using the CopyControl HTP Fosmid Library Production Kit (Epicentre) following the manufacturer's protocol. A library of ~ 4,000 clones were picked into 96-well plates and stored in –80 ºC as 50% glycerol stocks.

### 4. Feeding experiments with $^{15}NO_2^–$, $15NO_3^–$, and $^{15}$N-L-aspartic acid

*Streptomyces alanosinicus* ATCC 15710 was grown in 25 mL of fermentation medium in 250 mL baffled flasks while shaking at 30 ºC as described above. Stock solutions of [$^{15}$N]-calcium nitrate ($NaNO_3$), [$^{15}$N]-sodium nitrite ($NaNO_2$), and [$^{15}$N]-L-aspartic acid (Cambridge Isotope Laboratories) in water were sterilized by passing through a 0.22-μm filter membrane. After 16 h, each of these nitrogen sources was added to the fermentation cultures to a final concentration of 1 mM. After four more days of fermentation at 30 ºC, the presence of labeled and unlabeled **5** was determined by LC–HRMS after removing cell debris as described above.

**5. Identification of the putative L-alanosine (*ala*) biosynthetic gene cluster**

A BLAST search using *Staphylococcus aureus* L-diaminopropionic acid biosynthetic enzyme SbnA (annotation: 2,3-diaminopropionate biosynthesis protein SbnA, NCBI Accession: WP_000570808.1) was carried out using Geneious and revealed eight homologs in the *Streptomyces alanosinicus* ATCC 15710 genome (e-value < 1E-1). The same search using SbnB (annotation: N-[(2S)-2-amino-2-carboxyethyl]-L-glutamate dehydrogenase SbnB, NCBI Accession: WP_001078456.1) revealed four homologs in the S*. alanosinicus* ATCC 15710 genome (e-value < 1E-1). Only two hits from these BLAST searches were co-localized. Inspection of the genome neighborhood of each homolog based on the rationale described in the main text revealed the putative *ala* gene cluster. Annotations of the open reading frames (ORFs) in the *ala gene* cluster are found in Table S2.

**6. Cloning, overexpression, and purification of AlaB, AlaC, AlaL, AlaI, and AlaJ**

Expression plasmids were assembled from a PCR-amplified insert fragment of the gene of interest and a PCR-amplified vector fragment from the plasmid of interest using NEB Gibson Assembly. Inserts were amplified using primers designed with NEB Builder for Gibson assembly to overlap with PCR-linearized vectors (Table S3). PCR mixtures contained 2.8 µL of mQ water, 400 pg (1.7 µL) of template DNA, 2.0 µL of 10 mM forward primer, 2.0 µL of 10 mM reverse primer, 9.0 µL of PtSuperFi enhancer solution and 22.5 µL of 2x PtSuperFi Master Mix. Thermocycling was carried out in a MyCycler gradient cycler (Bio-Rad) using the following parameters: denaturation for 1 min at 98 ºC; 35 cycles of 0.16 min at 98 ºC, 1 min at 57 ºC, 2.5 min at 72 ºC; and a final extension time of 5 min at 72 ºC. PCR reactions were analyzed by agarose gel electrophoresis with ethidium bromide or SYBR safe staining, pooled, and purified. DNA was purified first by agarose gel electrophoresis and further purified with Zymoclean™ Gel DNA Recovery Kit. The PCR-amplified insert was ligated into the PCR-linearized expression vector using Gibson Assembly® Master Mix (New England Biolabs) following the manufacturer's instructions. 2 µL of each assembly reaction was transformed into a single tube of chemically competent *E. coli* TOP10 cells (Invitrogen). The identities of the resulting constructs were confirmed by sequencing of the purified plasmid DNA (Eton Bioscience). Successfully assembled plasmids were then transformed into chemically competent *E. coli* BL21 (DE3) cells (Invitrogen) and stored at –80 °C as frozen glycerol stocks.

**General Procedure for purification of His$_6$-tagged proteins.** Overnight starter cultures were inoculated from single colonies or from glycerol stocks made from single colonies. Cells were harvested by centrifugation at 4,000 rpm for 10 min, and the cell pellet was resuspended in 30 mL of lysis buffer (Buffer A: 20 mM HEPES, 500 mM NaCl, 10 mM $MgCl_2$, 10% *v/v* glycerol, pH 8.0). Cells were lysed via passage through a cell disruptor (Avestin EmulsiFlex-C3) twice at 10,000 psi, and the cell debris was pelleted via centrifugation at 10,800 rpm for 30 min. The supernatant was loaded onto Ni-NTA resin (1 mL, washed with Buffer A first) and washed with 30 mL of wash buffer (Buffer B: 25 mM imidazole, 20 mM HEPES, 500 mM NaCl, 10 mM $MgCl_2$, 10% *v/v* glycerol, pH 8.0). Bound protein was eluted with a minimal volume of elution buffer (Buffer C: 500 mM imidazole, 20 mM HEPES, 500 mM NaCl, 10 mM $MgCl_2$, 10% *v/v* glycerol, pH 8.0). Fractions containing desired protein were pooled, transferred to a Corning® Spin-X® UF 20 mL Centrifugal Concentrator (10,000 MWCO Membrane) and centrifuged at 4,000 rpm for 30 min to give ~200 µL of concentrate. The concentrate was diluted with 5 mL of Buffer A and centrifuged at 4,000 rpm for 30 min. The resulting concentrate was frozen in liquid nitrogen as ~20 μL beads and stored at –80 ºC. Individual protein beads were thawed before use and any excess was discarded to prevent repeated freeze-thaw cycles.

**General Procedure for purification of Strep-tagged proteins.** The same procedure was followed as described for the $His_6$-tagged proteins except that the clarified lysate was loaded onto Strep-Tactin Sepharose resin (1 mL, washed with Buffer A first) via gravity filtration. The resin was then washed with 3x 5mL of Buffer A. Bound protein was eluted with a minimal volume of elution buffer (Buffer D: Buffer A + 0.535 mg/mL of desthiobiotin).

**N-$His_6$-AlaB and N-$His_6$-AlaC.** 700 mL of sterile LB with ampicillin (100 μg/mL) in a 2.8 L Fleischman flask was inoculated with 2.5 mL of an overnight culture of *E. coli* BL21(DE3) containing the plasmids pETDuet_MCS1_alaB or pETDuet_MCS1_alaC. This was grown at 37 ºC and 180 rpm for 3 h or until an $OD_{600}$ of ~0.6 was reached, at which point IPTG was added to a final concentration of 100 μM. The culture was then returned to the incubator at 15 ºC and 180 rpm for 14 h after which AlaC was isolated following the general procedure for purification of $His_6$-tagged proteins as described above.

**N-$His_6$-AlaL-C-$His_6$.** 700 mL of sterile LB with kanamycin (50 μg/mL) in a 2.8 L Fleischman flask was inoculated with 20 mL of an overnight culture of *E. coli* BL21(DE3) containing the plasmid pET28a_alaL. This was grown at 37 ºC and 180 rpm for 1.5 h or until an $OD_{600}$ of ~0.6 was reached, at which point IPTG was added to a final concentration of 500 μM. The culture was then returned to the incubator at 15 ºC and 180 rpm for 14 h, after which AlaL was isolated following the general procedure for purification of $His_6$-tagged proteins as described above.

**N-$His_6$-AlaI.** 1 L of sterile LB with kanamycin (50 μg/mL) in a 2.8 L Fleischman flask was inoculated with 20 mL of an overnight culture of *E. coli* BL21(DE3) containing the plasmid pET28a_alaI. This was grown at 37 ºC and 180 rpm for 1.5 h or until an $OD_{600}$ of ~0.6 was reached, at which point IPTG was added to a final concentration of 100 μM. The culture was then returned to the incubator at 15 ºC and 180 rpm for 16 h, after which AlaI was isolated following the general procedure for purification of $His_6$-tagged proteins as described above.

**N-Strep-AlaJ.** 1 L of sterile LB with ampicillin (100 μg/mL) in a 2.8 L Fleischman flask was inoculated with 20 mL of an overnight culture of *E. coli* BL21(DE3) containing the plasmid pPR-IBA2-N-Strep_alaJ. This was grown at 37 ºC and 180 rpm for 1.5 h or until an $OD_{600}$ of ~0.6 was reached, at which point IPTG was added to a final concentration of 100 μM. The culture was then returned to the incubator at 15 ºC and 180 rpm for 16 h, after which AlaJ was isolated following the general procedure for purification of Strep-tagged proteins as described above.

**N-$His_6$-Sfp.** 1 L of sterile LB with ampicillin (100 μg/mL) in a 2.8 L Fleischman flask was inoculated with 20 mL of an overnight culture of *E. coli* BL21(DE3) containing the plasmid pET29b_Sfp. This was grown at 37 ºC and 180 rpm for 1.5 h or until an $OD_{600}$ of ~0.6 was reached, at which point IPTG was added to a final concentration of 800 μM. The culture was then returned to the incubator at 37 ºC and 180 rpm for 2 h, after which Sfp was isolated following the general procedure for purification of $His_6$-tagged proteins as described above.

### 7. *In vitro* biochemical assays

**NADH consumption assay of AlaB activity.** In a 50 µL reaction mixture in a 96-well plate (Greiner UV-Star®), 50 mM potassium phosphate pH 8.0, 10 mM $MgCl_2$, 2 mM L-Dap, 2 mM L-$\alpha$-ketoglutarate, and 50 µM of PLP were mixed. The reaction was initiated by the addition of 1 µM of AlaB. NADH consumption was monitored by measuring the decrease of absorbance at 340 nm. For results, see Figure S4.

**$^1$H NMR assay of AlaB activity.** In a 500 µL reaction mixture, 50 mM potassium phosphate pH 8.0, 10 mM $MgCl_2$, 2 mM L-Dap, 2 mM $\alpha$-ketoglutarate, 50 µM of PLP, and 10 µM of AlaB were mixed and incubated overnight at room temperature. The reaction mixtures were flash frozen with liquid $N_2$ and lyophilized. The residues were resuspended in $D_2O$ (Cambridge Isotope Inc.) and analyzed with proton nuclear magnetic resonance ($^1$H NMR) spectroscopy using an Agilent DD2-600 NMR spectrometer (600 MHz). Chemical shifts (δ) are reported in parts per million (ppm) downfield from tetramethylsilane using the solvent resonance as an internal standard for $^1$H NMR ($D_2O$ = 4.79 ppm). For results, see Figure S5.

**ATP-$^{32}$PPi exchange assay with AlaC.** To test the activity of the putative adenylation domain AlaC, 100 µL reaction mixtures were set up containing 50 mM HEPES (pH = 8), 200 mM NaCl, 10 mM $MgCl_2$, 5 mM dithiothreitol, 5 mM ATP, 1 mM of amino acid substrate, and 4 mM $Na_4PPi$/[$^{32}$P]PPi (3-5 x $10^5$ cpm/mL; Perkin Elmer). Each reaction was initiated via the addition of AlaC (1 µM) and incubated for 30 min at room temperature. Reaction mixtures were quenched with 200 µL of charcoal suspension (1.6% *w/v* activated charcoal in 100 mM $Na_4PPi$ with 3.5% *v/v* $HClO_4$), centrifuged (13,000 rpm for 3 min), and decanted. The charcoal pellet was washed (2x 200 µL) with wash buffer (100 mM $Na_4PPi$ with 3.5% *v/v* $HClO_4$). The charcoal pellet was then resuspended in 300 µL of wash buffer and added to 10 mL of scintillation fluid (Ultima Gold, Perkin Elmer) in a 25 mL scintillation vial. Radioactivity was subsequently measured on a Beckman LS 6000 scintillation counter.

**BODIPY-CoA assay to assess activity of AlaL.** A 50 µL reaction mixture was set up to test the ability of AlaL to be phosphopantethenylated. To a solution of reaction buffer (20 mM HEPES, 40 mM NaCl, 1 mM $MgCl_2$, pH = 8) was added BODIPY-coenzyme A[2] (5 µM), Sfp (1 µM), and AlaL (5 µM). This reaction mixture was allowed to incubate at room temperature for 2 h and was then combined with 50 µL of 2x SDS Laemmli sample buffer (BioRad) and boiled at 98 ºC for 15 min. Aliquots were loaded onto an SDS-Page gel and imaged at 365 nm and with Coomassie staining.

**Loading of L-Dap onto AlaL.** A 50 µL reaction mixture was set up to test the ability of AlaL to accept L-Dap. To a solution of reaction buffer (50 mM HEPES, 200 mM NaCl, 10 mM $MgCl_2$, pH = 8) was added coenzyme A (1 mM), Sfp (5 µM), and AlaL (20 µM). This reaction was allowed to incubate at room temperature for 2 h, then AlaC (20 µM) and L-2,3-diamonpropionic acid or L-$^{15}N_2$-Dap (250 µM) were added followed by ATP (5 mM) to initiate the reaction. After incubating for 1 h at room temperature, reactions were halted by flash-freezing in liquid nitrogen. Samples were thawed, centrifuged at 13,000 rpm in a table-top centrifuge and submitted for analysis.

High-resolution LC-MS analyses were performed in the Small Molecule Mass Spectrometry Facility at Harvard University on an Agilent 6220 TOF Mass Spectrometer fitted with an electrospray ionization (ESI) source. Liquid chromatography was performed on an Agilent Technologies 1200 series LC using Agilent PLRP-S polymeric column (50 x 2.1 mm, 1000 Å pore size, 5 µm packing material; injection volume 5 µL) at a flow rate of 0.25 mL/min and the following elution conditions using 0.1% formic acid in water as mobile phase A and 0.1% formic acid in acetonitrile as mobile phase B. The column temperature was maintained at room temperature. The following gradient was applied: 0-10 min: 5- 50% B in solvent A, 10-25 min: 50% B isocratic, 25-26 min: 50-5% B, 26-31 min: 5% B isocratic. Elution before 2 min was diverted to waste. All experiments were performed in the positive ionization mode for a mass range of m/z 400 to 3000 with the m/z scale externally calibrated using Agilent ESI-L Low Concentration Tuning Mix (P/N: G1969-85000). Deconvolution of protein raw mass spectra was performed using Agilent MassHunter software (version B.06.00). The masses detected corresponded to $[M+H]^+$ ions and are listed in Table S5.

**8. Gene disruption experiments**

Gene inactivation in *Streptomyces alanosinicus* ATCC 15710 was performed according to standard protocols.[3] Briefly, primers used to amplify *alaD* were used to screen the fosmid library to obtain a fosmid containing the *ala* gene cluster. The fosmid was transformed into *E. coli* BW25113/pkD46 by electroporation. The *aac(3)IV*-oriT cassette was amplified by PCR from pIJ773 using the primers listed in Table S4. Each of the biosynthetic genes was replaced with the *aac(3)IV*-oriT cassette PCR product using PCR targeting and λ-red-mediated recombination. The mutant fosmid was transformed into *E. coli* WM6026 for conjugation with *Streptomyces alanosinicus* ATCC 15710. Double crossover mutants were selected with apramycin resistance on MS agar, and the exconjugants were grown in TSB with apramycin added (50 µg/mL final concentration). After the seed culture reached saturation, the genomic DNA was isolated, and insertion of the resistance marker was confirmed by PCR. The mutants were tested for production of **5** under the same fermentation conditions as described in Section 2 for the wild-type strain with the exception of added antibiotics.

For the chemical complementation experiments, wild-type and mutant strains were grown in 10 mL of fermentation medium in 50 mL baffled flasks while shaking at 30 ºC as described in section 2. Stock solutions of $NO_2^-$ in water were sterilized by passing through a 0.22 µm filter membrane and added to the fermentation to 1 mM final concentration after 16 h. After 2 more days of fermentation at 30 ºC, the presence of **5** was determined by LC-HRMS as described in Section 2.

**9. Griess assay for nitrite (**$NO_2^-$ production)

An 80 µL reaction mixture was set up to test the ability of N-$His_6$-AlaI and N-Strep-AlaJ to generate $NO_2^-$ from L-aspartic acid. To a solution of reaction buffer (50 mM HEPES, 10 mM $MgCl_2$, pH = 8.0) was added FAD (10 µM), NADPH (5 mM), N-Strep-AlaJ (5 µM), and N-$His_6$-AlaI (5 µM). Reactions were initiated by the addition of L-aspartic acid (1 mM) and incubated at room temperature for 1 h. Reactions were quenched via the addition of 80 µL of MeOH, vortexed briefly, and centrifuged at 13,000 rpm for 10 min to pellet the precipitated protein. A 30 µL aliquot of the supernatant was combined with 30 µL of 1.0% sulfanilic acid in 1 N HCl and 30 µL of 0.2% (napthyl)ethylenediamine dihydrochloride in 1 N HCl. The resulting mixture was transferred to a 96 well plate and the absorbance at 548 nm was recorded with a BioTek Gen5 Microplate Reader.

### 10. Chemical synthesis and compound characterization

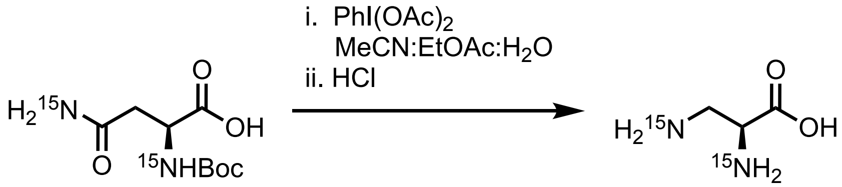

A 2 dram vial containing a stir bar and Boc-L-Asn-OH (52.6 mg, 0.226 mmol) was charged with water (0.3 mL), MeCN (0.6 mL), and EtOAc (0.6 mL). The resulting clear, colorless solution was cooled to ~0 ºC in an ice/water bath. To this was added PIDA (87.2 mg, 0.271 mmol) in one portion. After 15 h, the stir bar was removed and the reaction mixture was concentrated *in vacuo* to give a white solid. The resulting solid was sonicated with ~1 mL of EtOAc, collected via vacuum filtration, and dried under house vacuum. The resulting off-white solid was dissolved in water (0.5 mL) and cooled to ~0 ºC with vigorous stirring. The pH of this solution was adjusted to ~ 1 via the addition of concentrated HCl, which resulted in the evolution of gas. This mixture was concentrated *in vacuo* and the crude yellow oil was purified using an aminopropyl Sep-Pak (Waters) and an elution gradient of 0 to 20 to 40 to 60 to 80 to 100% water in acetonitrile. Product-containing fractions (determined by TLC in 3:1:1 *n*-BuOH:water:AcOH and stained with ninhydrin) were combined and lyophilized to give a L-Dap•HCl as a white solid (3.6 mg, 0.0252 mmol, 11.1% yield over 2 steps).

For $^{15}N_2$-L-Dap, HRMS (ESI+) Calcd. for $C_3H_9{}^{15}N_2O_2$ : 107.0599 Found: 107.0577. $^1H$ NMR data matched previous reports.[4]

**Data Availability**

The nucleotide sequences for the *ala* biosynthetic gene cluster and individual genes have been deposited into NCBI (accession numbers: TBD). Additional data that support the conclusions of the paper can be requested from the corresponding author.

**Figure S1.** SbnA and SbnB produce L-Dap from *O*-phospho-L-serine and L-glutamate.[5]

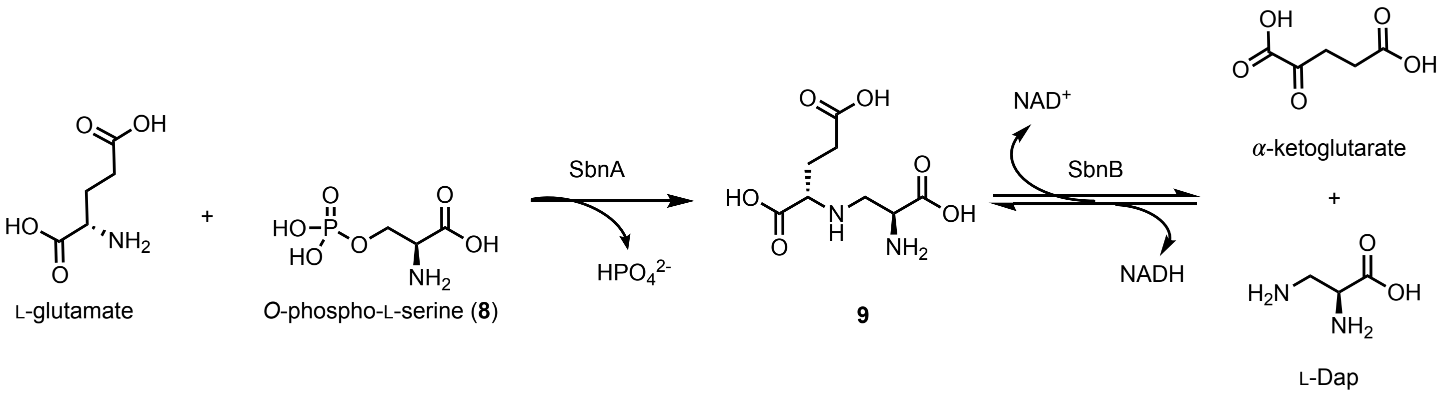

**Figure S2.** LC-MS/MS analysis of $^{15}$N-**5** from feeding of $^{15}NO_2^-$

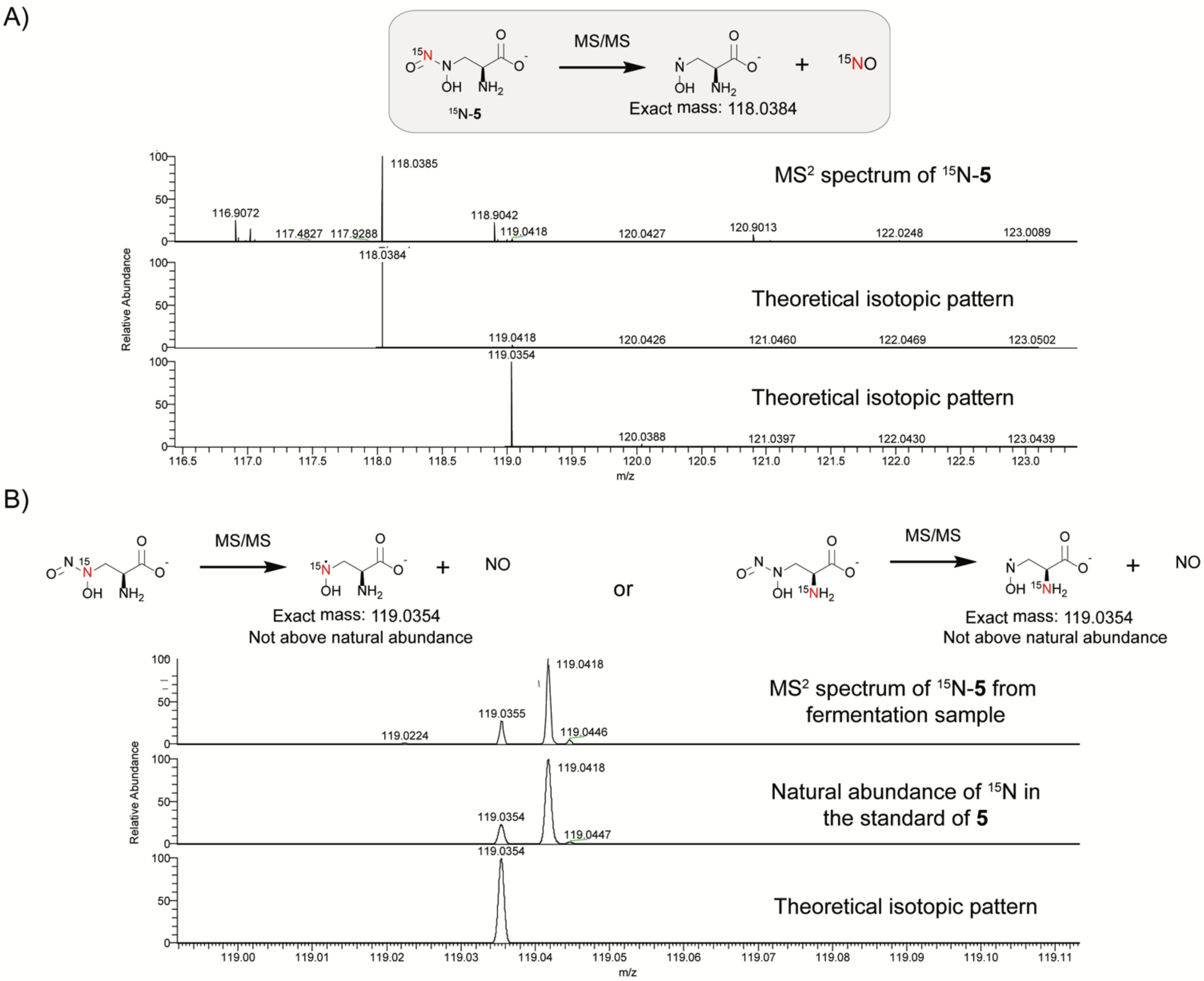

To determine which nitrogen atom of **5** was enriched, we performed LC-MSMS analysis. If the distal *N*-nitroso nitrogen is labeled, we would expect an unlabeled hydroxylamine radical fragment ion (panel a). If the nitrogen in the proximal *N*-nitroso nitrogen or the amino group of **5** are $^{15}$N-labeled, we would expect enrichment of $^{15}$N in the hydroxylamine radical fragment (panel b). We observed an unlabeled hydroxylamine radical fragment with no enrichment of $^{15}$N above natural abundance indicating that the distal *N*-nitroso nitrogen is labeled.

**Figure S3.** Mass spectrum of **5** from cultures fed with 3 mM of $^{15}NO_2^-$

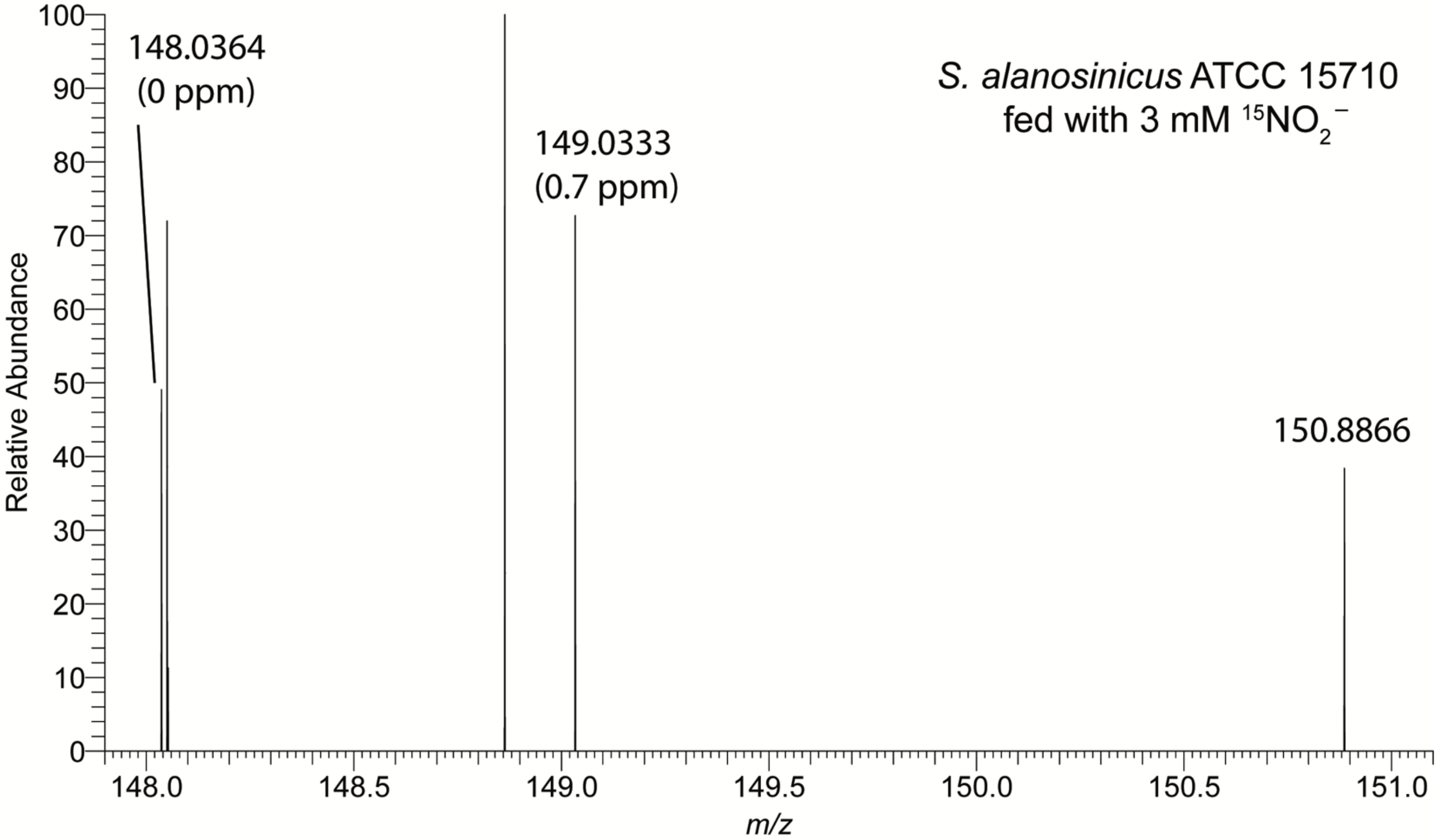

The exact [M–H]$^-$ mass of **5** and $^{15}$N-**5** are 148.0364 and 149.0334 respectively.

**Figure S4.** Mass spectrum of **5** from cultures fed with $^{15}NO_3^-$

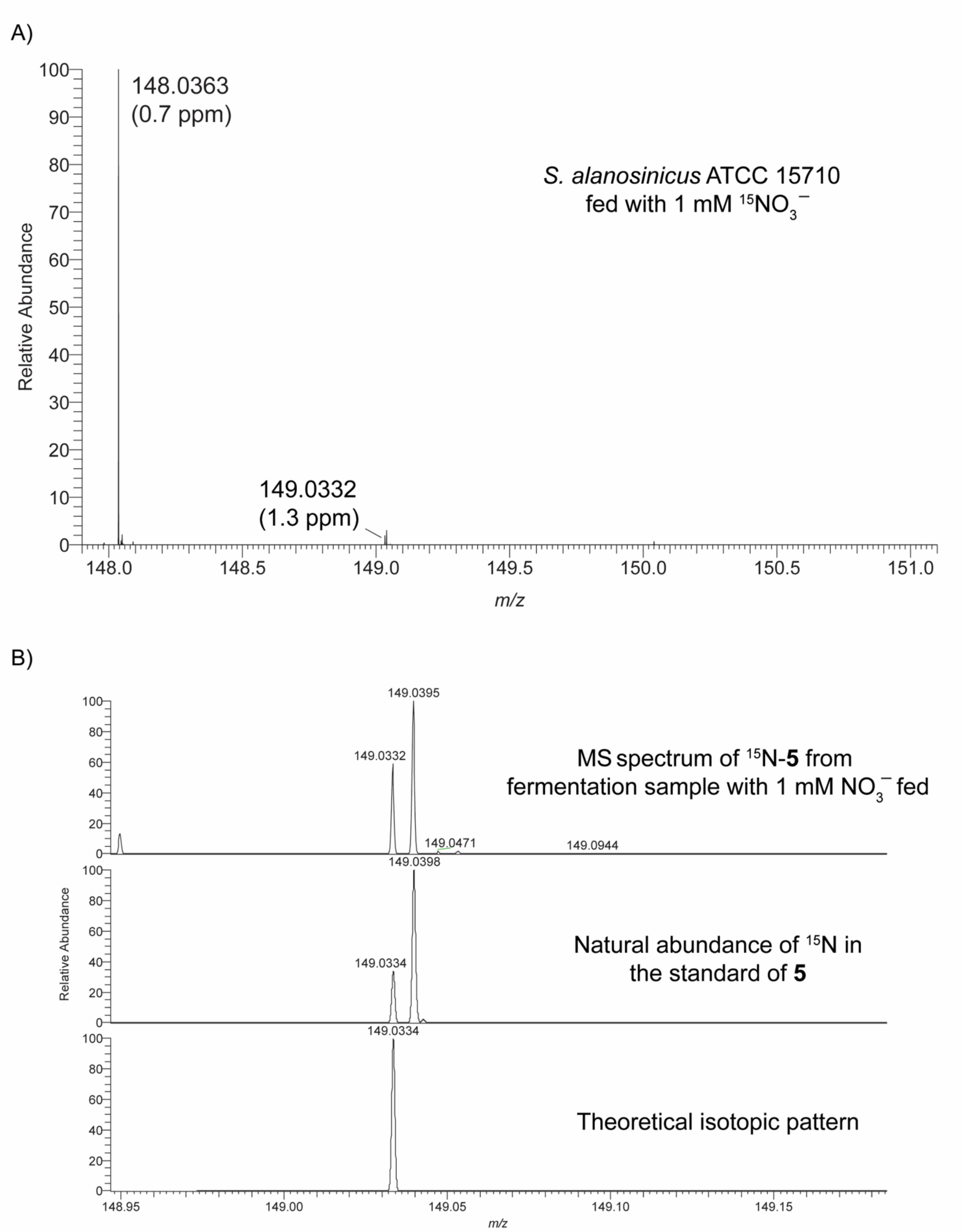

The exact [M–H]$^-$ mass of **5** and $^{15}$N-**5** are 148.0364 and 149.0334 respectively.

**Figure S5.** AlaB consumption of NADH with L-Dap and α-ketoglutarate

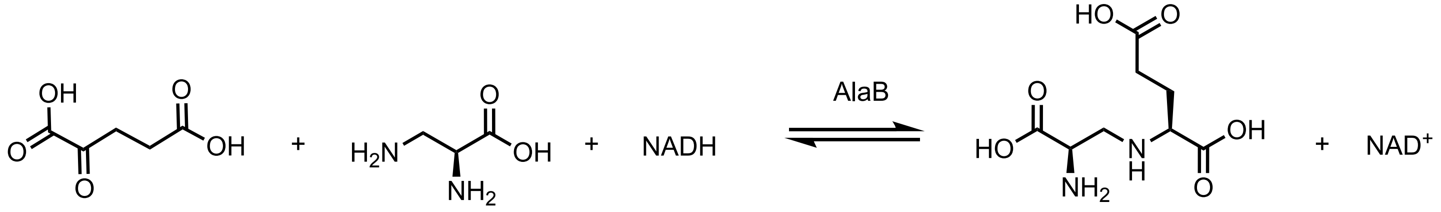

α-ketoglutarate L-Dap **9**

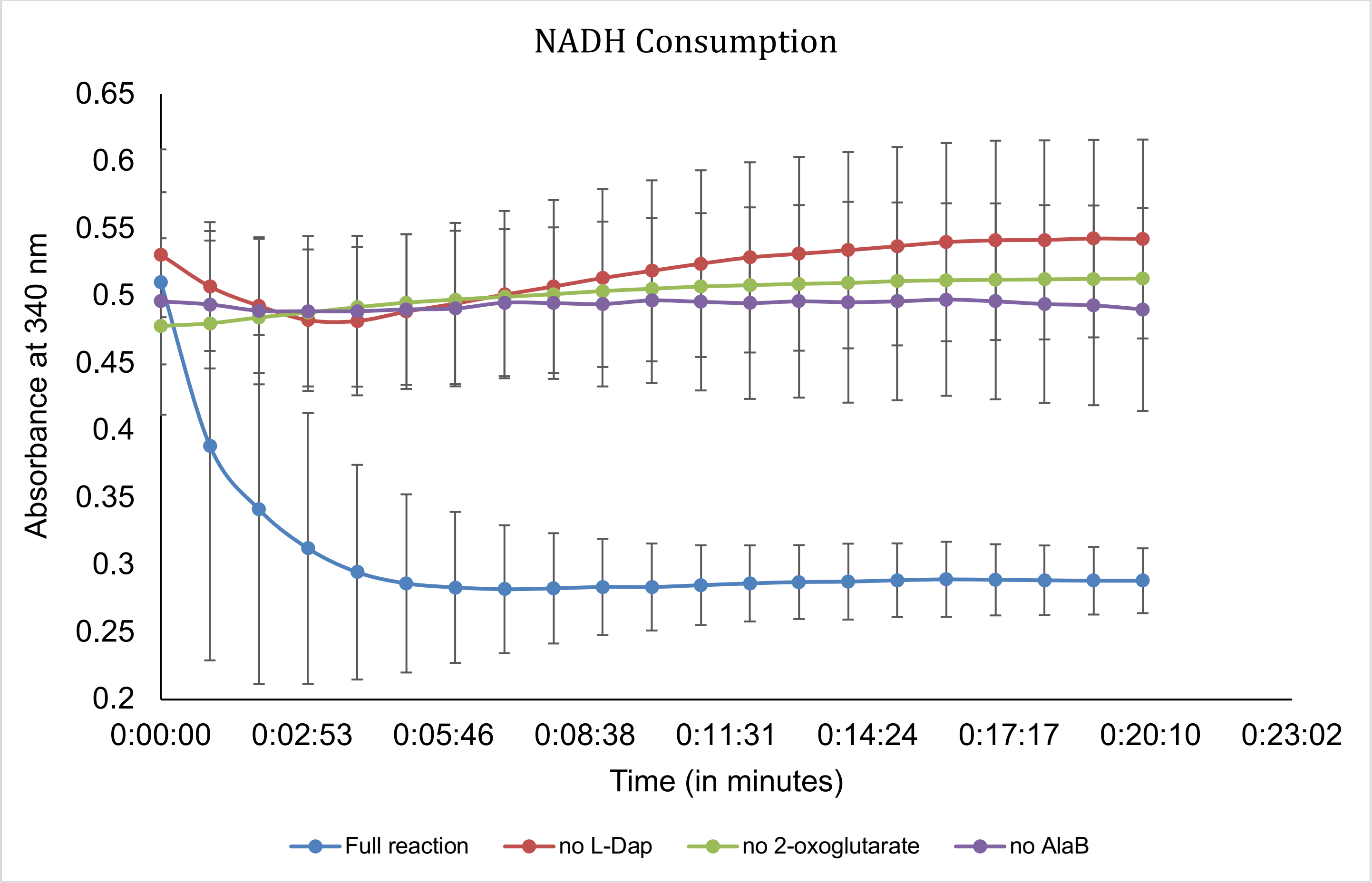

Each reaction was performed in triplicate. One set of technical replicates was performed on a different day. Data are mean ± standard deviation (s.d.) of three biological replicates. The assay conditions are reported in the Methods section above.

**Figure S6.** $^{1}$H NMR of **9** from *in vitro* biochemical assay of AlaB with L-Dap, α-ketoglutarate, and NADH

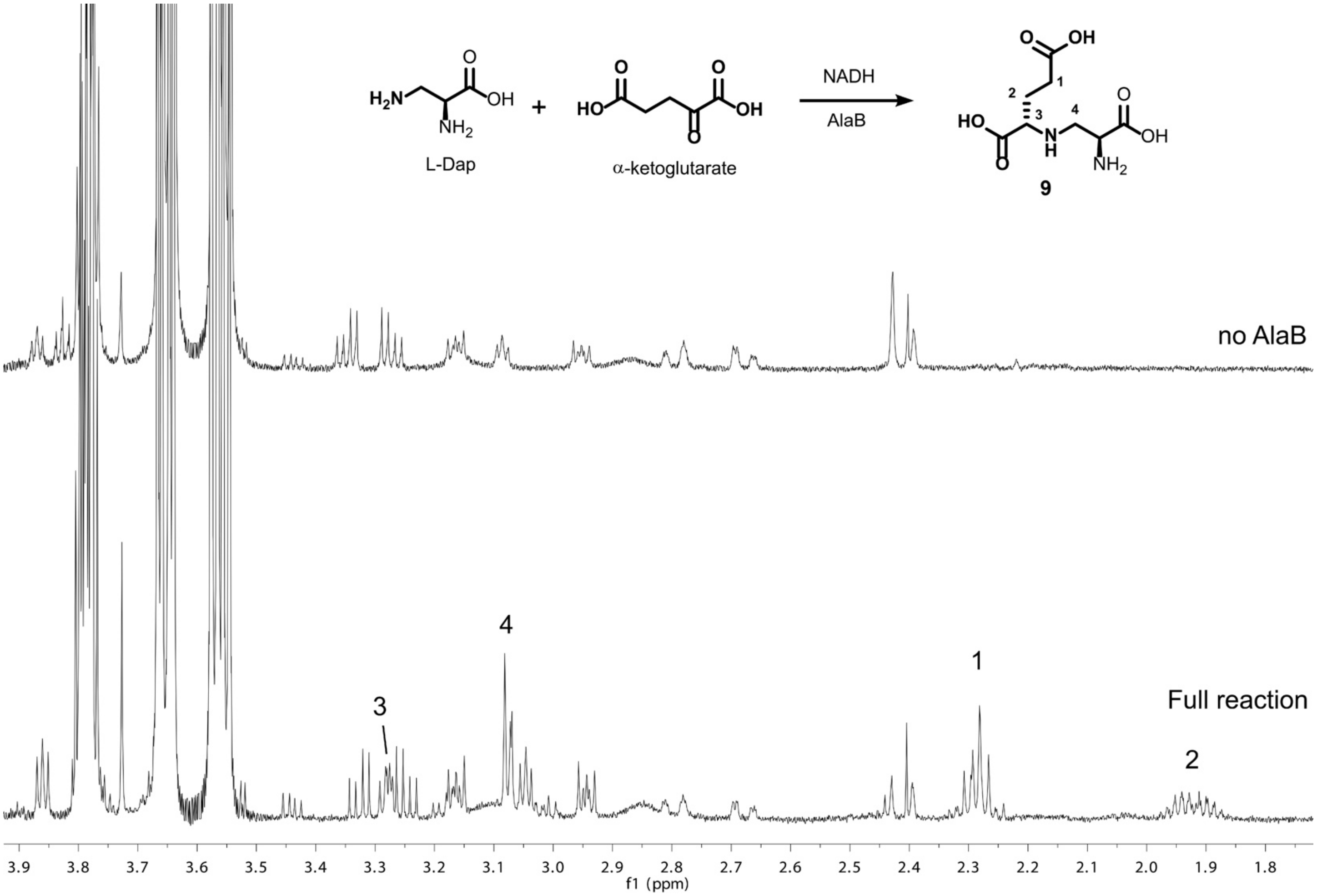

Top spectrum: A reaction with L-Dap, α-ketoglutarate, and NADH without AlaB added. Bottom spectrum: The full reaction in which L-Dap, α-ketoglutarate, AlaB, and NADH are added. The expected chemical shifts for product **9** were assigned according to published literature.[6]

**Table S1.** Annotations of the *ala* biosynthetic genes.

| Protein | Size (aa) | Predicted Function | Closest Homolog | Accession # | %Identity/ %Similarity |
|---|---|---|---|---|---|
| AlaA | 324 | 2,3-diaminopropionate biosynthesis protein | SbnA, *Streptomyces hirsutus* | WP_055594931.1 | 87/91 |
| AlaB | 353 | 2,3-diaminopropionate biosynthesis protein | SbnB, *Streptomyces hirsutus* | WP_055594930.1 | 84/89 |
| AlaC | 557 | amino acid adenylation domain-containing protein | *Streptomyces hirsutus* | WP_055594929.1 | 79/84 |
| AlaD | 385 | acyl-CoA dehydrogenase | *Streptomyces hirsutus* | WP_055594947.1 | 89/94 |
| AlaE | 190 | hypothetical protein | *Streptomyces hirsutus* | WP_055594928.1 | 80/88 |
| AlaF | 237 | thioesterase | hypothetical protein, *Streptomyces hirsutus* | WP_079034952.1 | 76/83 |
| AlaG | 286 | hypothetical protein | *Streptomyces hirsutus* | WP_055594925.1 | 86/92 |
| AlaH | 177 | NADPH-dependent FMN reductase | *Streptomyces hirsutus* | WP_055594946.1 | 87/92 |
| AlaI | 657 | FAD/NAD(P)-binding protein | FAD/NAD(P)-binding protein [*Streptomyces kanamyceticus*] | WP_055544225.1 | 76/82 |
| AlaJ | 470 | nitro-succinate lyase | 3-carboxy-*cis,cis*-muconate cycloisomerase [*Streptomyces sp. BK161*] | TDT22211.1 | 75/80 |
| AlaK | 273 | SARP family regulator | CreF [*Streptomyces cremeus*] | ALA99205.1 | 57/69 |

**Table S2.** Analysis of adenylation domain specificity for AlaC and other putative L-Dap activating A domains.

| Organism | *Streptomyces alanosinicus* | *Streptomyces albulus PD-1* | *Bacillus thuringiensis* | *Streptomyces vinaceus* |
|---|---|---|---|---|
| **Natural Product** | L-alanosine (**1**) | poly(L-DAP) | zwittermicin A | viomycin |
| **A domain** | AlaC | EXU85975.1 | ZWAA2 | VioA1 |
| **Accession #** | N/A | EXU85975.1 | ACD36028.1 | AAP92491.1 |
| **A domain "codon"** | D I W E L T A D | D F E Y V G T V | D V F Y L G G V | D V Y H F S L V |
| **Maryland NRPS-PKS Prediction**[7] | No hit | Pip | Pps3-M2-Val | Cda1-M1-Ser |
| **SEQL-NRPS Prediction**[8] | TYR | DHPG-DPG | PRO | SER |
| **NRPSsp Prediction**[9] | PHE | PHE | ornithine | serine |
| **Experimentally Confirmed** | L-Dap† | L-Dap† | L-Dap‡ | L-Dap† |
| **Reference** | - | 10 | 11 | 12 |

† measured via ATP-PPi exchange; ‡ measured via EnzCheck Pyrophosphate Assay Kit

**Table S3**. Primers used for cloning for protein expression.

| Oligo | Nucleotide Sequence (5' to 3') | Targeted gene |
|---|---|---|
| AlaB-F | GCCTGCAGGTCGACAAGCTTAGCGATCAGGCCAGCATCCC | *alaB* |
| AlaB-R | CTTAAGCATTATGCGGCCGCTCGCTGCCCCCTCTCCGG | *alaB* |
| AlaC-F | GCCTGCAGGTCGACAAGCTTATGACGACCGTCCCTTCTG | *alaC* |
| AlaC-R | CTTAAGCATTATGCGGCCGCCGAGGCCTCATTCCGGTG | *alaC* |
| AlaL-F | TGCCGCGCGGCAGCCATATGATGACGGCGACCTGGCAG | *alaL* |
| AlaL-R | TCGAGTGCGGCCGCAAGCTTTGGGGTGAGAAAATTCTGCTCG | *alaL* |
| AlaI-F | ATACATATGGAGAGAGGGACTATGAC | *alaI* |
| AlaI-R | ATAAAGCTTTCATCGCGTCAGTTCTCCGA | *alaI* |
| AlaJ-F | GTTCGAAAAAGGCGCCAGCATCATCGATGCCGGGCTG | *alaJ* |
| AlaJ-R | CAAGCTTAGTTAGATATCTTATGCGTCCGCCTCGCGCAG | *alaJ* |

**Table S4**. Primers used for generating knockout cassettes from pIJ773.

| Oligo | Nucleotide Sequence (5' to 3') | Targeted gene |
|---|---|---|
| AlaC-KO-F | CATGTCCGTGCACACGACAACGCGACCGCCCTGCTCGAC-ATTCCGGGGATCCGTCGACC | *alaC* |
| AlaC-KO-R | GGGCACCATCGCGGTCGGCAGTGCCGTGCTCAGCGCGTC-TGTAGGCTGGAGCTGCTTC | *alaC* |
| AlaD-KO-F | GCTCTCGCGGAGATGCGCCGCACCGGTCTGCTCGGGCTG-ATTCCGGGGATCCGTCGACC | *alaD* |
| AlaD-KO-R | GGTCTTTTCCAGCCGGGTAGGCGAGTTCTGGAGATAGCC-TGTAGGCTGGAGCTGCTTC | *alaD* |
| AlaI-KO-F | GTGTCGACGCCCGACGACGTGGCGATCACCGTGCATGTGAT-TCCGGGGATCCGTCGACC | *alaI* |
| AlaI-KO-R | CACCGAGCCGACTCCCGGGCGGATGCCGGCCGCCGTCGC-TGTAGGCTGGAGCTGCTTC | *alaI* |

**Table S5.** N-His$_6$-AlaL-C-His$_6$ masses detected corresponding to $[M+H]^+$ ions with loss of the terminal methionine residue and loading of L-Dap or L-$^{15}N_2$-L-Dap.

| Species | Formula | Calculated Mass (Da)* | –Met | Detected Mass (Da) $[+H]^+$ |
|---|---|---|---|---|
| apo-AlaL | $C_{575}H_{887}N_{169}O_{175}S_6$ | 13151.4028 | 13028.56 | -- |
| holo-AlaL | $+C_{11}H_{21}N_2O_6PS$ | 13499.85 | 13368.65 | 13369.60 |
| L-Dap-AlaL | $+C_{14}H_{26}N_4O_7PS$ | 13585.89 | 13454.69 | 13455.97 |
| L-$^{15}N_2$-Dap-AlaL | $+C_{14}H_{26}N_4O_7PS$ | +1.9942 | 13456.68 | 13457.69 |

*mass calculated from the amino acid sequence for N-His$_6$-AlaL-C-His$_6$: MGSSHHHHHHSSGLVPRGSHMMTATWQDVRSWILQRNPELAEDELNPETDIIESRIVDSLQFVELVL FIEELRGEMMQSMDVDLDSFRTLKDIEQNFLTPKLAAALEHHHHHH
